# Cell-free DNA profiling of metastatic prostate cancer reveals microsatellite instability, structural rearrangements and clonal hematopoiesis

**DOI:** 10.1101/319855

**Authors:** Markus Mayrhofer, Bram De Laere, Tom Whitington, Peter Van Oyen, Christophe Ghysel, Jozef Ampe, Piet Ost, Wim Demey, Lucien Hoekx, Dirk Schrijvers, Barbara Brouwers, Willem Lybaert, Els Everaert, Daan De Maeseneer, Michiel Strijbos, Alain Bols, Karen Fransis, Steffi Oeyen, Pieter-Jan van Dam, Gert Van den Eynden, Annemie Rutten, Markus Aly, Tobias Nordström, Steven Van Laere, Mattias Rantalainen, Prabhakar Rajan, Lars Egevad, Anders Ullén, Jeffrey Yachnin, Luc Dirix, Henrik Grönberg, Johan Lindberg

## Abstract

**Background:** There are multiple existing and emerging therapeutic avenues for metastatic prostate cancer, with a common denominator, which is the need for predictive biomarkers. Circulating tumor DNA (ctDNA) has the potential to cost-efficiently accelerate precision medicine trials to improve clinical efficacy and diminish costs and toxicity. However, comprehensive ctDNA profiling in metastatic prostate cancer to date has been limited.

**Methods:** A combination of targeted- and low-pass whole genome sequencing was performed on plasma cell-free DNA and matched white blood cell germline DNA in 364 blood samples from 217 metastatic prostate cancer patients.

**Results:** ctDNA was detected in 85.9% of baseline samples, correlated to line of therapy and was mirrored by circulating tumor cell enumeration of synchronous blood samples. Comprehensive profiling of the androgen receptor (AR) revealed a continuous increase in the fraction of patients with intra-*AR* structural variation, from 15.4% during first line mCRPC therapy to 45.2% in fourth line, indicating a continuous evolution of AR during the course of the disease. Patients displayed frequent alterations in DNA repair deficiency genes (18.0%). Additionally, the microsatellite instability phenotype was identified in 3.81% of eligible samples (≥0.1 ctDNA fraction). Sequencing of non-repetitive intronic- and exonic regions of *PTEN, RB1* and *TP53* detected biallelic inactivation in 47.5%, 20.3% and 44.1% of samples with ≥0.2 ctDNA fraction, respectively. Only one patient carried a clonal high-impact variant without a detectable second hit. Intronic high-impact structural variation was twice as common as exonic mutations in *PTEN* and RB1. Finally, 14.6% of patients presented false positive variants due to clonal hematopoiesis, commonly ignored in commercially available assays.

**Conclusions:** ctDNA profiles appear to mirror the genomic landscape of metastatic prostate cancer tissue and may cost-efficiently provide somatic information in clinical trials designed to identify predictive biomarkers. However, intronic sequencing of the interrogated tumor suppressors challenge the ubiquitous focus on coding regions and is vital, together with profiling of synchronous white blood cells, to minimize erroneous assignments which in turn may confound results and impede true associations in clinical trials.

## Background

Prostate cancer is the most commonly detected male cancer in Europe and the third major cause of cancer-related death among men [1]. Although the majority of metastatic hormone-naïve prostate cancers (mHNPCs) demonstrate a reliable response to initial androgen deprivation therapy which targets AR signaling, progression to a castrate resistant state is inevitable. However, the treatment landscape for metastatic castrate resistant prostate cancer (mCRPC) is evolving with the recent approval of several new drugs translating to an increased overall survival [2–6]. Multiple additional avenues exist as genomic profiling of metastatic tissue revealed that the majority of mCRPC patients harbor clinically relevant alterations beyond the AR signaling pathway [7].

The most promising non-approved treatment avenue in metastatic prostate cancers (mPC) exploits synthetic lethality in treating homologous recombination deficient cancers with poly (ADP-ribose) polymerase (PARP) inhibitors [8]. Approximately one fifth of mCRPC carry mutations in DNA repair genes [7]. However, the mutational signatures of biallelic inactivation is heterogeneous between different DNA-repair genes [9] and future studies are therefore needed to determine which genes are associated with a response to PARP inhibition. Approximately three percent of mPC are driven by microsatellite instability (MSI) [7,10]. Pembrolizumab recently became the first drug to be approved by the U.S. Food and Drug Administration based on the MSI phenotype, irrespective of tumor type [11]. Although check-point blockade did not confer a survival advantage as compared with placebo for chemotherapy-relapsed mCRPC [12], anecdotal cases have been reported to display partial or complete responses [10,13–15].

The emergence of additional drugs, both towards common and rare mPC phenotypes such as PTEN-deficient [16,17] and neuroendocrine cancers [18], raises questions of how to efficiently translate the multitude of treatment options to improved patient outcomes. The genomic heterogeneity of mCRPC [7] and in turn, the low response rates of currently approved drugs [2–5,19,20], argue for the urgent need of predictive biomarkers. Ineffective trial-and-error decisions inevitably lead to unnecessary side-effects and unsustainable costs [21]. The AR splice variant 7 (AR-V7) [22] demonstrated promising results as a negative response biomarker for abiraterone and enzalutamide [23]. However, follow-up studies did not validate the initial clear-cut finding and the clinical value of AR-V7-assays remain debated [24].

The lack of predictive biomarkers is in part due to the difficulty of obtaining temporally matched metastatic tissue as the majority of mPCs metastasise to the bone. Multiple studies on the acquisition of tumor tissue with or without direct image-guidance report a range of success rates [25–28]. A recent effort, focusing on methodological improvements, obtained >20% cell content in the majority of bone biopsies [29]. Circulating tumor DNA is a viable alternative to metastatic tissue with demonstrated high fractions of ctDNA [30–33] enabling sensitive detection of somatic variation and direct comparisons to metastatic tissue reveal high concordance [30,34,35]. Circulating tumor DNA has several advantages as sampling through simple blood draws is fast, cost efficient, without side-effects, allows for longitudinal monitoring and the detection of multiple resistance alleles during therapy [35,36].

Although ctDNA has the potential to accelerate biomarker-driven trials in mPC several questions remain unanswered, e.g. if it is possible to detect MSI directly from liquid biopsies and how ctDNA fractions correlate to line of therapy. The ctDNA fraction determines the sensitivity to detect somatic variation which in turn has consequences for the design of prospective biomarker studies relying on liquid biopsies. Here we present a retrospective analysis of 217 cases and 364 blood samples covering the entire spectrum of mPC. The purpose of this study was to gather information relevant for future liquid biopsy-driven biomarker studies with a focus on 1) how ctDNA fractions vary from mHNPC to end stage castration-resistant disease; 2) a rationale for how to treat samples with low ctDNA fraction; 3) the relative impact of different types of somatic variation, affecting the sequencing strategy; 4) the detection of potentially predictive biomarkers; 5) and finally, how clonal expansions in the hematopoietic stem cells [37–40] impact liquid biopsy profiling.

## Methods

### Patient cohorts

The ProBio (Prostate Biomarkers) cohort was collected in the Stockholm area during 2015–2017 with the goal to perform retrospective hypothesis-generating liquid biopsy analysis for future studies. Metastatic prostate cancer patients ranging from hormone-naïve to castration-resistant were invited to participate and asked to donate blood. Anonymous healthy donor blood was collected from healthy men at Hötorgets Blood Centre in central Stockholm. Additionally, mCRPC cases were recruited in Belgium via a non-interventional clinical study (CORE-ARV-CTC study) between March 2014 and April 2017. The purpose of this cohort study was to investigate if profiling androgen receptor splice variants in circulating tumor cells (CTCs) may predict enzalutamide and abiraterone treatment response. Leftover plasma was biobanked simultaneously and used for ctDNA profiling, described here. Accounting for both cohorts, cell-free DNA and germline DNA profiling was performed on 217 mPC and 36 healthy donors (Table 1). In addition, the FDA-cleared CellSearch CTC technology (Menarini Silicon Biosystems, Italy) was applied for CTC enumeration on 340 out of 364 blood samples processed for ctDNA analysis. Ethical approval was obtained from ethical committees in Belgium (Antwerp University Hospital, registration number: B300201524217) and Sweden (Stockholm Regional Ethical Vetting Board registration numbers: 2016/101-32, amendment 2017/252-32; 2009/780-31/4, amendment 2014/1564-32). All patients provided a written informed consent document.

**Table 1.**
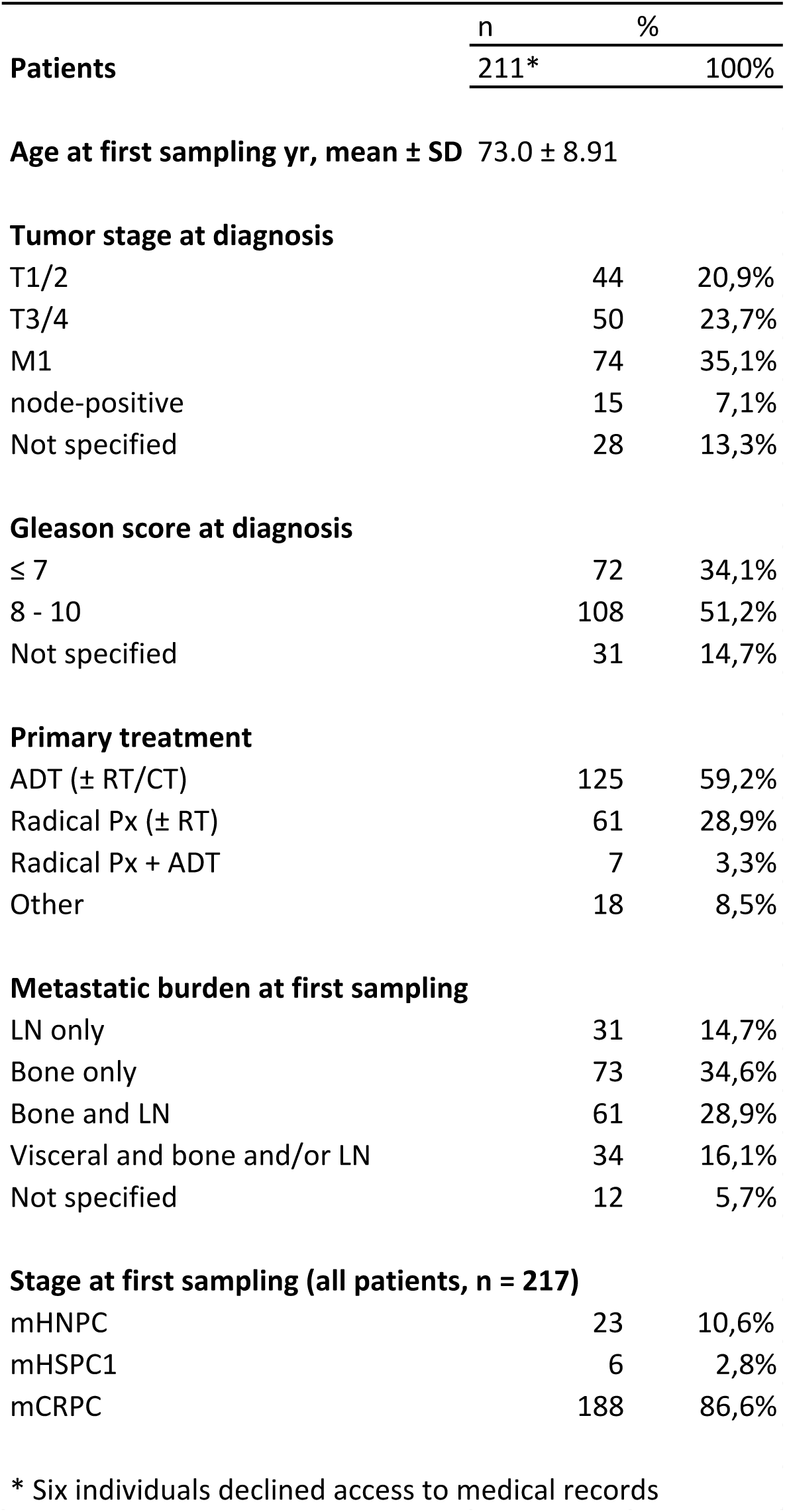
Clinical characteristics describing the study participants.

### Sample processing and sequencing

In ProBio, plasma was enriched from 2x10 ml EDTA blood whereas 4–5 ml of blood was available from the CORE-ARV-CTC study. Germline DNA was extracted from leftover EDTA blood after plasma centrifugation. In both studies, the EDTA blood tubes were processed within the same working day, allowing for high-quality ctDNA profiling [41]. Blood collected in CellSave tubes were shipped to the GZA Sint-Augustinus hospital in Antwerp, to perform CTC counting within 72 hours, as previously described [42]. The plasma was stored at −80°C until cell-free DNA (cfDNA) isolation. cfDNA was isolated using the QiaSymphony system (Qiagen, Germany). Purified cfDNA was subjected to fragment analysis for quality control. Germline DNA from WBCs was isolated using the AllPrep DNA/RNA Micro Kit (Qiagen, Germany). Library prep was mainly performed using the ThruPLEX Plasma-seq kit (Rubicon Genomics, USA). 0.1 – 50 ng of cfDNA and 50 ng of germline DNA was used to create the sequencing libraries. Cell-free DNA profiling was performed with a mix of low-pass whole genome sequencing and targeted sequencing allowing for identification of copy-number alterations (CNAs), small mutations and structural variation. The SeqCap EZ system (Roche Nimblegen, USA) was applied for targeted sequencing. The targeted regions were designed to capture unique regions in the human genome commonly mutated in prostate cancer, identified through literature review. The designs are described in Supplementary table 1. Briefly, the comprehensive designs were aimed at progression samples, targeting common single nucleotide polymorphisms at a certain intervals to enable CNA detection. The smaller designs were tailored for cost efficient deep sequencing in combination with low-pass whole genome sequencing for profiling CNAs. Follow-up samples and samples with <5 ng of cfDNA available were profiled with a smaller, focused design, to enable cost-efficient saturation of the library. Low-pass whole genome sequencing was applied to baseline samples with <5 ng cfDNA to enable CNA analysis.

### Sequence data analysis

Low-level processing of the sequencing data was performed as previously described [33]. Statistical analysis and filtering of variants was conducted in R [43]. Twenty-five plasma samples displayed significantly increased fraction of discordantly mapped read pairs and low allele frequency variants. Five had previously been profiled from the same blood draw without any quality issues [33]. To avoid false positives, these samples were not included for structural variation- or MSI analysis. In addition, mutations below five percent allele fraction were discarded unless they were detected in another sample of the same individual.

The data from our previous publications [33,41] was merged with samples that were processed again to increase coverage. Only non-default settings are displayed for the analysis tools below. Germline small variants in blood samples were identified with FreeBayes [44] (version: 1.0.1, settings: --min-alternate-fraction 0.01 --min-coverage 20, then filtered on QUAL > 5) and annotated with VEP [45] (version: 83, settings: --pick --filter_common --check_alleles -- check_existing --total_length --allele_number --no_escape --no_stats --everything --offline). Heterozygous SNPs were identified and their allele ratio in the cfDNA samples were used in analysis of copy number alteration and loss of heterozygosity. Downstream analysis of putative oncogenic germline variants was performed in R. Variants were required to be supported by ≥12 reads and ≥20% allele ratio, and to be either annotated as *pathogenic* or *likely pathogenic* in ClinVar [46], or introduce a premature stop or frameshift in a coding sequence. Findings were inspected manually and annotated for evidence of somatic LOH based on the cfDNA copy number profiles and their allele ratio of heterozygous SNPs.

Somatic point mutations in cfDNA samples were identified with VarDict [47] (settings: -f 0.01 -Q 10, followed by var2vcf_paired.pl with settings -P 0.9 -m 4.25 -M -f 0.01 keeping variants flagged as LikelySomatic or StrongSomatic), using patient-matched blood samples as controls and annotated with VEP (same settings as above). Downstream analysis was performed in R. Known hotspot mutations in *AKT1, APC, AR, ATM, BRAF, CDKN2A, CTNNB1, DICER1, DNAJB1, EGFR, ERBB2, FOXA1, GNAQ, HOXB2, HRAS, IDH1, IDH2, KAT8, KDM6A, KRAS, LRP1, MAP2K1, MED12, PDGFRA, PIK3CA, PIK3CB, PIK3R1, PPP2R1A, PTEN, RAF1, SMAD4, SPOP, TP53* and *XPO1* were not filtered further (≥1% allele frequency). Remaining variants were required to be supported by ≥6 reads and ≥2% allele ratio, and to not be observed ≥2 times in the set of healthy donor cfDNA samples (Supplementary figure 1). Variants without any effect on coding sequence were also removed except for purity estimation. Remaining variants were reviewed manually and removed if reoccurring in multiple patients at low allele ratio without being a known hotspot mutation. Germline DNA processing failed for four patients (P-GZA3843, P-D6, P-AZSJ044 P-AZSJ055). Healthy donor cfDNA was used as reference to identify point mutations in these patients retaining only high-impact or hotspot mutations. Variants were also annotated for evidence of somatic LOH based on the cfDNA copy number profiles and their allele ratio of heterozygous SNPs. Sublonal variants were pragmatically defined as having an allele frequency < 1/4 of the ctDNA fraction, a similar approach as previously applied for ctDNA in colorectal cancer [48].

Normalized coverage (log_2_-ratio) and segmentation for copy number analysis was produced with qDNAseq [49] for low-pass WGS (version: 1.8.0, settings: binSize=15, only using forward reads) and with CNVkit [50] for targeted sequence data (version: 0.7.9). The allele ratio of heterozygous SNPs was used with the log-ratio and segmentation to verify and curate observations. Putative somatic focal amplification was called where the median log-ratio at the gene exceeded control regions to the left and right (3–8Mb from gene start or end) by at least 0.5. Putative somatic focal deletion was similarly called where the log-ratio of both control regions exceeded that of the gene by at least 0.3. Amplifications and deletions were curated manually and considered real if supported by the SNP allele ratio and if not attributable to poor data quality such as low coverage and waviness [51]. Deletions were considered homozygous if no large genomic segments (≥5Mb) had a similar or lower median log-ratio, if the SNP allele ratio of the segment did not indicate allelic imbalance, and if adjacent segments indicated hemizygous deletion or LOH. The sensitivity to detect homozygous deletions and LOH was compromised below 0.2 ctDNA fraction but is also affected by sequence coverage and the clonality of the individual event. Therefore, only samples with ctDNA fraction ≥ 0.2 were used to investigate if one high-impact variant could suffice to infer biallelic inactivation. Somatic deletions indicating *TMPRSS2-ERG* fusion were identified manually from log-ratio and SNP allele ratio. Germline copy number deletion was called where segmented log-ratio of the normal DNA was below −0.5.

The svcaller algorithm was applied for the identification of structural variants as described previously [33]. To reduce the proportion of false positive calls the following filters were implemented:

1. Filter out a read pair if there exists an alternative mapping (of equal mapping quality) that is consistent with normal genomic positioning (i.e. pointing towards each other, with an insert size < 1000bp). This eliminates a class of visually apparent false positive calls caused by mapping issues.
2. Reject an event if more than half of the reads have identical start position at either of the termini. This excludes a class of visually apparent false-positive events, in particular frequent weakly supported translocation calls.
3. Filter based on soft-clipped read clustering for duplications or inversions. Exclude the event if the event contains scattered soft-clip regions. An event is considered to have scattered soft-clip regions if the ratio of grouped soft-clips to the number of groups is <= 4. Soft-clippings are first assigned to groups by considering the start and end position of each soft-clippied read and subsequently to the group with the largest soft-clipping count.
4. Filter out events that have overlapping, shared termini: Retain the single largest event, when an overlap occurs.

To improve performance two initial filters were applied:

1. Filter out events where all contributing reads have a mapping quality of zero.
2. Only consider read pairs where both reads map to chromosomes 1–22, X and Y.

Manual evaluation was performed on each variant candidate. Structural variants of unknown significance that occured in multiple germline samples were filtered out. A sample was considered to have a significant structural variants in *AR* if it contained any genomic structural rearrangement that: 1) did not affect exons upstream of cryptic exon 4; 2) affected any exon, including- and downstream of cryptic exon 4. However, non-functional variation such as tandem duplications with the 5’ breakpoint in the ligand binding domain and the 3’ breakpoint downstream of AR, were excluded. In addition, structural events removing *AR* exon 1, known to lead to AR45 expression [52], were considered to be significant structural variants. All other events in *AR* were considered structural variants of unknown significance. Subclonal point mutations were defined as having an allele frequency < 1/4 of the ctDNA fraction. However, comparing the allele frequencies of structural variants *(PTEN, TMPRSS2-ERG* region, *TP53, RB1)* to commonly mutated genes in mPC *(AKT1, APC, ATM, BRAF, BRCA1, BRCA2, CDK12, CHEK2, CTNNB1, FOXA1, KRAS, PIK3CA, PTEN, RB1, SPOP, TP53)*, in samples that harbored both, revealed consistently lower structural variant allele frequencies (Supplementary figure 2). The ratio between the median structural variant allele frequency and the median mutation allele frequency was 0.275. Therefore, a structural variant was considered to be subclonal if the allele frequency was < 0.0688 (0.275*0.25) of the ctDNA fraction. After applying this threshold for structural variants on all samples, the number of subclonal structural variants in *ERG, PTEN, TMPRSS2, TP53* and *RB1* was not statistically different from mutations (Number subclonal and clonal for: structural variants, 24 and 162; mutations, 79 and 387. x2 test: p = 0.201). The current version of the svcaller software is available online: https://github.com/tomwhi/svcaller.

Tumor burden (ctDNA component in cfDNA) was initially estimated from the median allele frequency of somatic point mutations with an allele frequency of at least 5%. However, the variant with highest allele frequency was copy-number adjusted and compared to the median allele frequency. If discordant (+/− 0.05) the ctDNA fraction was corrected. Where somatic copy number alterations suggested a higher tumor burden, it was instead calculated from the difference in log-ratio associated with gain or loss of one copy. Only low-fraction structural variation was detected in for P-00030277_3042897, P-00041183_3094920, P-00043867_3124162 and P-NIKO005_20170077. For these four, the average of the number of supporting reads divided by the total read depth at the two breakpoint positions was used as ctDNA fraction estimate.

Evaluation of the sensitivity and specificity to detect MSI was performed on an in-house cohort of 450 colorectal cancer samples out of which 61 had MSI. Microsatellite instability was evaluated by 1) applying the mSINGS [53] algorithm 2) comparing the number of mutations, separated into single nucleotide variation and indels. A more stringent threshold was applied (≥10 supporting reads) to filter out artefact variants as intronic and synonymous variants were included. 3) manual inspection of copy-number alteration patterns, where microsatellite unstable tumors have limited copy-number alterations unlike the chromosomal instability phenotype. The MSI colorectal tumors, were diluted in silico to determine a purity- and a fraction unstable microsatellites cutoff for the mSINGS algorithm. Applied on the whole set, a purity cutoff of 10% and and mSINGS 0.1 fraction of unstable microsatellites demonstrated 100% sensitivity and 99% specificity.

Mosaic copy number alterations in blood DNA (smaller effect than consistent with ±1 copy per cell) were considered indications of clonal hematopoiesis. Additional small variants indicating clonal hematopoiesis were identified with VarDict, using patient blood samples with a merged file of all healthy donor blood samples as a control, and annotated with VEP. They were validated in matched cfDNA with VarDict (patient cfDNA with unmatched healthy donor as control). Known hotspot mutations were not filtered further. Other variants supported by ≥6 reads and an allele ratio of 2–25%, with effect on coding sequence, without indication of being SNPs or mapping errors (manual curation) were considered indicative of clonal hematopoiesis.

## Results

### Liquid biopsy profiling of metastatic prostate cancer

Comprehensive cell-free DNA (cfDNA) profiling was performed on 217 mPC patients (Table 1). Single nucleotide variants, copy-number alterations (CNAs) and genomic structural rearrangements were interrogated using a combination of in-solution hybridization capture-based and low-pass whole genome sequencing. An evolution of capture designs was applied as the project progressed, from a pancancer to a prostate-specific approach to cost-efficiently maximize the information content (Supplementary table 1). The comprehensive designs were aimed at progression samples with high tumor burden whereas the smaller designs were tailored for cost-efficient deep sequencing. However, to increase the sensitivity to detect e.g. intronic structural variation in *AR*, the majority of samples were processed with both a comprehensive- and a smaller design (Supplementary table 2). The data was subsequently merged before variant calling. The average coverage, taking merging into account, was 814x (interquartile range: 251 - 965) for cfDNA, and 445x (interquartile range: 371 - 533) for germline DNA. Data from all samples is presented here, where the number of relevant samples is described for each section. In total 364 plasma samples from 217 mPC patients were profiled. Circulating tumor cell (CTC) enumeration using the CellSearch platform was performed from synchronous blood draws on 340 of the 364 plasma samples.

### Circulating tumor DNA fraction correlation to line of therapy

Circulating tumor DNA was detected in the majority of baseline samples (85.9%). However, the fraction of ctDNA in the cfDNA determines the sensitivity to detect somatic variation. We therefore investigated if tumor burden correlated to blood draw timing and line of therapy (Figure 1, Supplementary table 3). The ctDNA fractions were significantly lower, comparing baseline and follow-up samples for mHNPC and first line mCRPC. The differences were not statistically significant at later lines of therapy. The CTC counts were correlated to the ctDNA fraction estimate (rho = 0.7, p < 0.0001) (Supplementary figure 3), and mimicked the ctDNA pattern in relation line of therapy (Figure 1). In addition, comparing baseline ctDNA fractions at different lines of therapy, a significant increase was only present between first to second line mCRPC and third to fourth line mCRPC (Supplementary figure 4).

**Figure 1.**
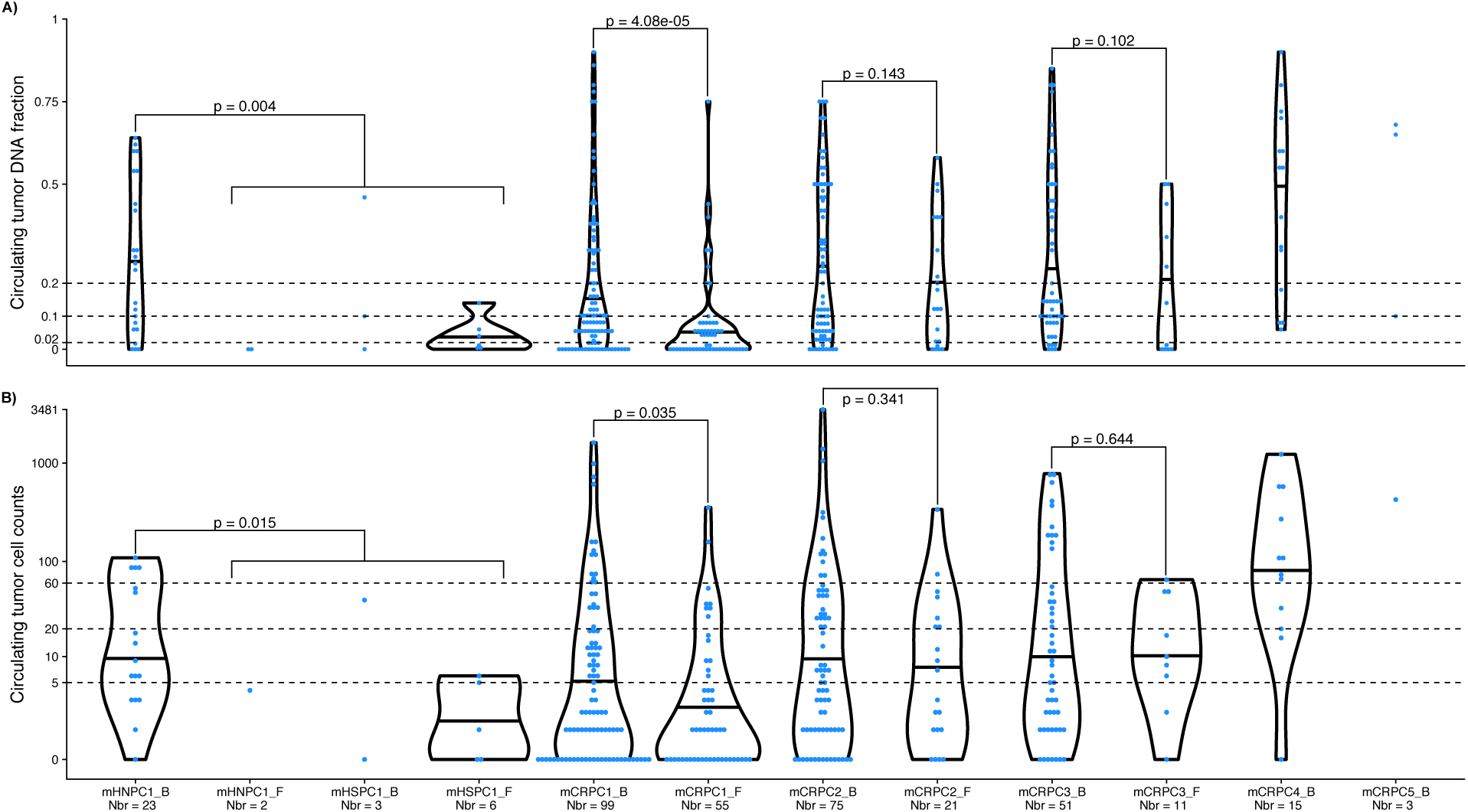
Tumor burden at different lines of therapy. **A)** Violin plot of the circulating tumor DNA fraction partitioned according to line of therapy. Black horizontal lines within the violin plots denotes the median circulating tumor DNA fraction. Blue points represent the circulating tumor DNA fraction in individual blood samples. A one-sided Wilcoxon rank sum test was applied to investigate if the baseline samples had higher tumor burden than the follow-up samples. Y-axis: Circulating tumor DNA fraction. X-axis: Line of therapy. **B)** as A) but for circulating tumor cell counts per 7.5 ml of blood. Y-axis: log10 transformed circulating tumor cell counts. Abbreviations: mHNPC[number], metastatic hormone naive prostate cancer and line of therapy; mHSPC[number], metastatic hormone sensitive prostate cancer and line of therapy; mCRPC[number], metastatic castrate resistant prostate cancer and line of therapy; _B, baseline; _F, follow-up; Nbr, number of samples profiled in each category.

### Detection of microsatellite instability from cell-free DNA

Microsatellites were targeted and sequenced to enable MSI-phenotype detection directly from cfDNA (Supplementary table 1). Using an in-house cohort of colorectal samples (Supplementary figure 5), in-silico dilution with germline DNA demonstrated 100% sensitivity and 99% specificity to detect MSI at 10% tumor purity with the mSINGS algorithm [53]. Applying mSINGS on ≥10% ctDNA fraction samples identified four cases with MSI out of 105 investigated (Figure 2). The proportion of MSI-positive cases detected from ctDNA is concordant with a previous study, performing whole-exome sequencing on metastatic tissue (Fisher's exact test: p = 0.721) [7].

**Figure 2.**
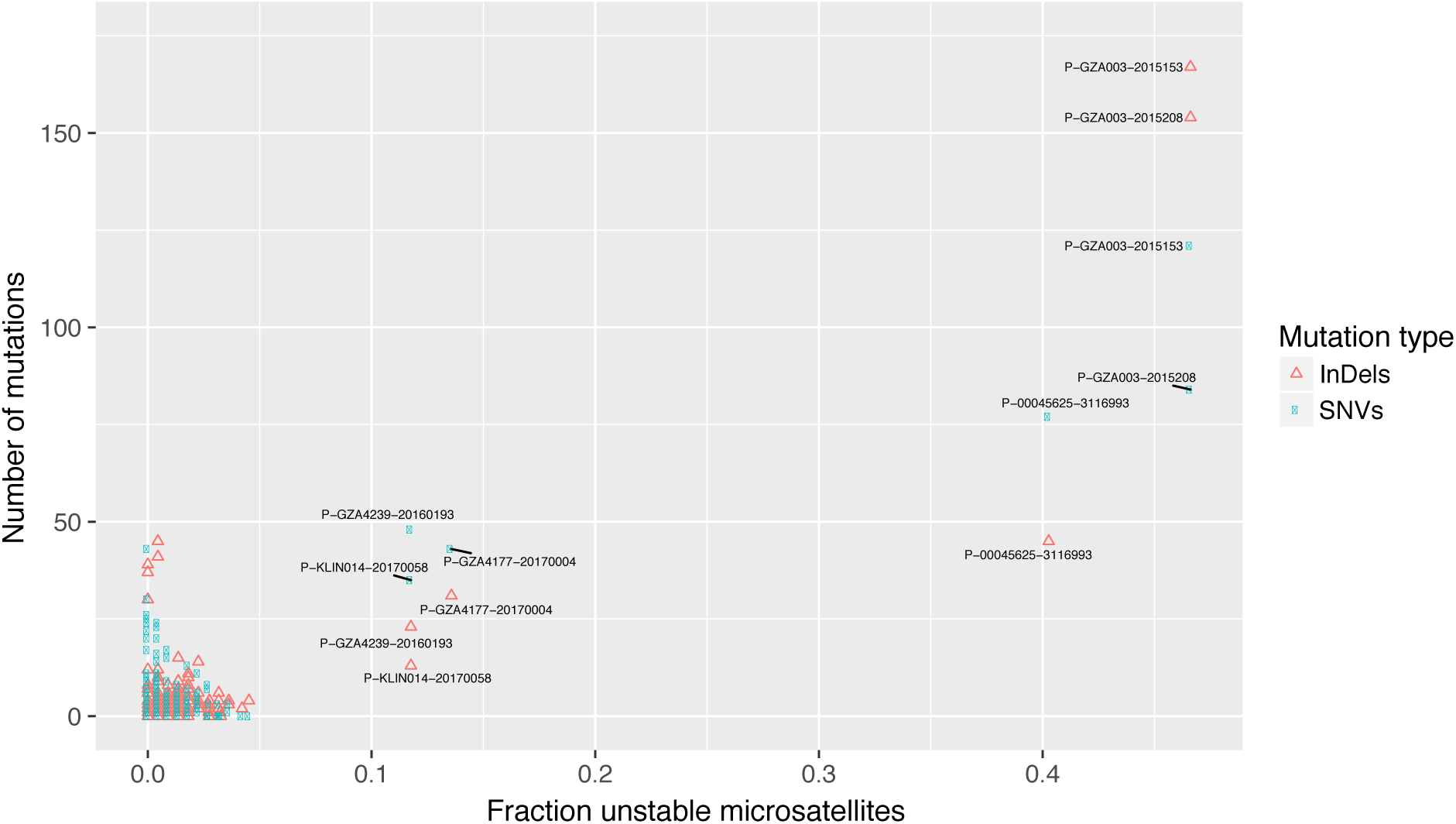
Detection of microsatellite instability from cell-free DNA. Microsatellite unstable tumors were identified by plotting the number of mutations (y-axis, including intronic and synonymous variants) versus the fraction of unstable microsatellite loci (x-axis). Indels and single nucleotide variants are kept separate for each sample, colored according to the right legend. Note, although individual P–KLIN014, sample 20170058 demonstrated >0.1 fraction unstable microsatellite loci, it was classified as microsatellite stable. The sample had high circulating tumor DNA fraction (0.8), lacked increase in mutational burden and displayed massive copy-number alterations, indicative of a chromosomal instability phenotype.

### Intronic sequencing of key tumor suppressors and biallelic inactivation

Prostate cancer is mainly driven by CNAs, commonly generated through chained structural rearrangements. Chained events cause the majority of *TMPRSS2-ERG* gene fusions [54], also observed in our data (Figure 3, Supplementary figure 6). To investigate the impact of structural rearrangements capture probes were designed towards the non-repetitive intronic and exonic regions of *PTEN, RB1* and *TP53* in the prostate specific comprehensive design (CP design, Supplementary table 1 and 4, Supplementary figure 7). Structural rearrangements, mutations and copy-number alterations were investigated in 165 cfDNA samples from 135 study participants profiled with the CP design that passed the internal quality control for structural variant calling (Methods). Seventy-one samples (71/165, 43.0%) from 59 men (59/135, 43.7%) had a ctDNA fraction ≥0.2, where all classes of somatic variation were detectable. Biallelic inactivation, through clonal high-impact variation, occurred in 47.5% (28/59), 20.3% (12/59) and 44.1% (26/59) of patients in *PTEN, RB1* and *TP53*, respectively. After excluding the MSI samples (carrying high-impact passenger mutations in multiple genes), all samples with a clonal high-impact variant also harbored a second event with only one exception: two samples were profiled for patient P-00030277 and both revealed a 392 kb deletion encompassing exon 9–10 of *TP53* without any observable second hit.

**Figure 3.**
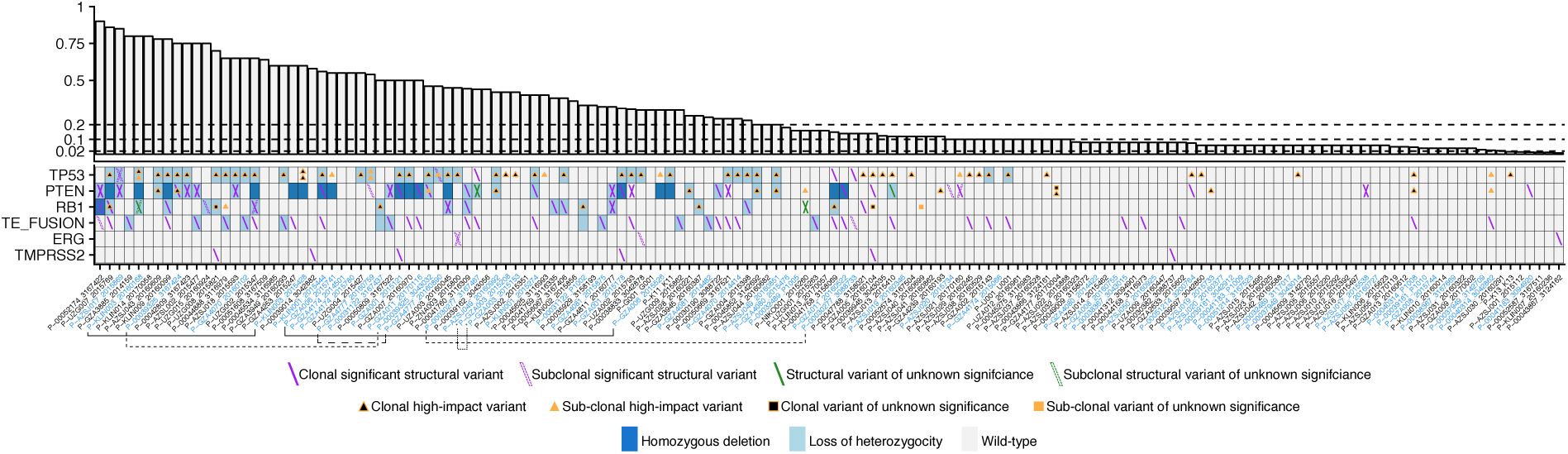
- Exonic and intronic profiling of circulating tumor DNA.

The non-repetitive sequence was captured for the entire gene body of *TP53, PTEN* and *RB1*. The samples with *TMPRSS2-ERG* gene fusions or structural rearrangements in *TMPRSS2* or *ERG* separately are also shown here. The upper panel displays the circulating tumor DNA fraction. The dashed lines at 0.02, 0.10 and 0.20 denotes the cutoff to detect point mutations, loss of heterozygosity and homozygous deletions, respectively. Bottom panel, heatmap of the mutational landscape detected from circulating tumor DNA profiling. Type of alteration is coded according to the bottom legend. For visualization purposes, maximally two mutations or structural variants are displayed per patient. Triangles and boxes represents single nucleotide variants and indels. Subclonal mutations are defined as having an allele frequency < 1/4 of the ctDNA fraction. The same definition was applied to structural variants after median allele allele frequency adjustment versus the mutations (Methods). Synonymous point mutations are not displayed here. Variants of unknown significance are non-synonymous single nucleotide variants outside hotspots, not annotated as pathogenic from variant databases. Structural variants of unknown significance are e.g. confined to a single intron, without affecting neighbouring exons. X-axis: Cell-free DNA samples sorted according to the circulating tumor DNA fraction. Patients with multiple samples are colored in blue. *samples with microsatellite instability. Samples described in the main text are connected with lines.

### Clonal dynamics during treatment

Subclonal high-impact variation was detected in multiple patients. Both samples of P-GZA006 revealed subclonal *TP53* mutation accompanied by subclonal deletion. Before start of abiraterone therapy, sample 20160759 of patient P-GZA4777 carried two subclonal *TP53* mutations (hotspot and frameshift), a subclonal translocation in *PTEN* and a weak *AR* amplification (Supplementary figure 8A). At progression (sample 20160890), the hotspot *TP53* variant, *PTEN* translocation and the *AR* amplification were undetectable. However, the *TP53* frameshift increased in allele fraction and a new structural deletion in *TP53* was found, in line with *TP53* loss being associated with rapid progression [55]. P-KLIN003 also experienced change in clonal composition during abiraterone therapy (Supplementary figure 8B). At baseline, two *TP53* mutations were detected. After therapy, the two displayed different behaviour, decreasing and increasing in allele fraction. The progressing clone also presented with *TP53* loss of heterozygosity (LOH) and multiple structural variants in AR. Patient P-00039325 had high ctDNA fraction despite being on androgen deprivation therapy for three weeks. Following docetaxel treatment, P–00039325 progressed after 215 days with a translocation in *BRCA2* and concomitant LOH (Supplementary figure 8C). In addition, an *AR* amplification and intra*-AR* structural variation were detected.

### Continuous evolution of somatic variation in the androgen receptor

Comprehensive profiling of *AR*, including intronic sequencing, was performed in 275 mCRPC plasma samples from 177 individuals (Figure 4A). In total 45.8% (126/275) of the samples and 50.3% (89/177) of the patients harbored one or more variant in *AR* (high-impact mutation, structural variant or amplification) in at least one cfDNA sample (Supplementary table 4). Intra-AR structural variation was closely correlated to amplification and only 3/51 patients (P-GZA4045, P-GZA4120, P-U001) carried intra-AR structural variation without an accompanying amplification. Structural variation was detected in another three patient samples without amplification (P-AZSJ022, P-KLIN002, P-UZA002), but weak amplifications were found in other samples from the same individuals, taken at other occasions. The fraction of patients with structural variation in *AR* correlated to line of therapy, ranging from 15.4% during first line mCRPC therapy to 45.2% in fourth line. Overall, the percentage of individuals with any alteration in *AR* increased from 37.4% in first line to 76.9 % in fourth line, indicating a continuous evolution of *AR* during the course of the disease (Figure 4B).

**Figure 4.**
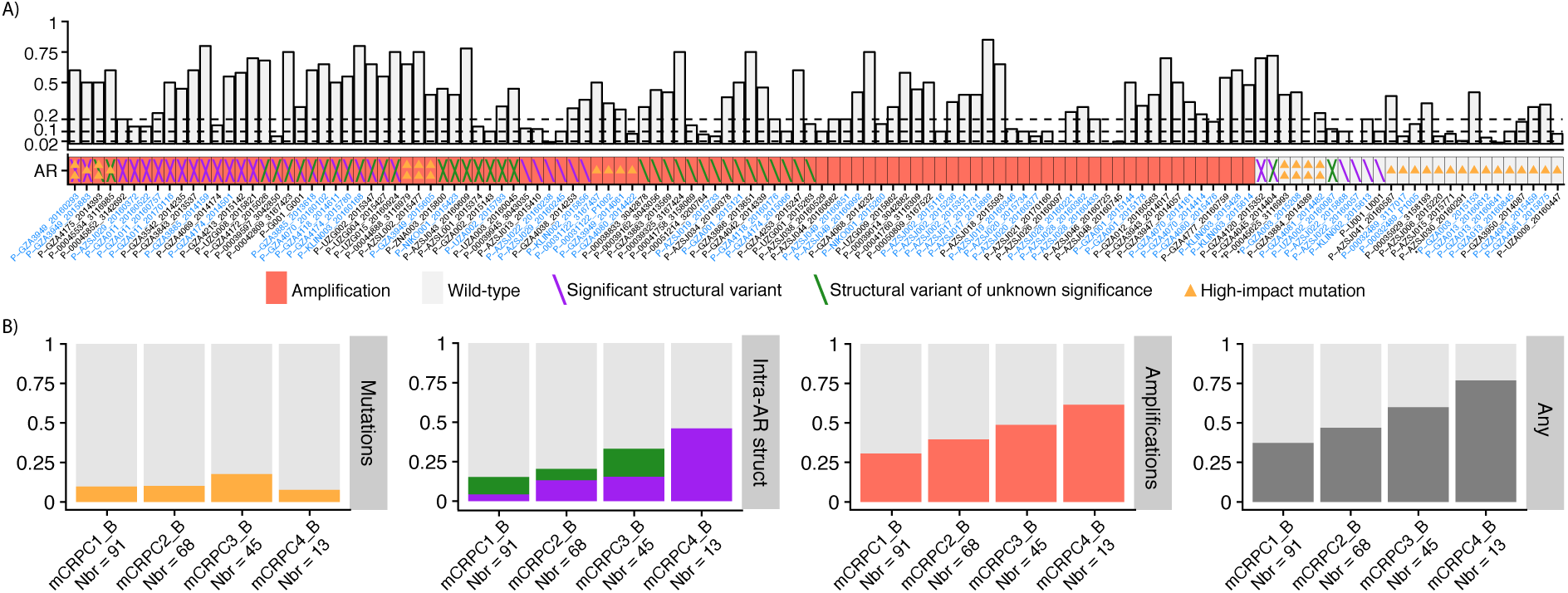
Androgen receptor alterations. **A)** The upper panel displays the circulating tumor DNA fraction. The dashed lines at 0.02, 0.10 and 0.20 denotes the cutoff to detect point mutations, loss of heterozygosity and homozygous deletions, respectively. Bottom panel, heatmap of the mutational landscape detected in the androgen receptor from circulating tumor DNA profiling. Type of alteration is coded according to the bottom legend. For visualization purposes, only samples with an alteration are shown here. Maximally two mutations or structural variants are displayed per sample. X-axis: Cell-free DNA samples sorted according to the number of alterations detected. Patients with multiple samples are colored in blue. *samples with microsatellite instability. **B)** The fraction of patients with alterations in the androgen receptor is categorized by type of alteration and line of therapy. Only high-impact mutations, e.g. hotspot mutations are accounted for here. Intra-AR structural variation is colored according to the legend in A) The rightmost bar plot represents the fraction of patients with alteration in the androgen receptor taking any type of variation into account. Abbreviations: mCRPC[number], metastatic castrate resistant prostate cancer an line of therapy; _B, baseline; Nbr, number of samples profiled.

### Alterations in DNA repair deficiency genes

Genes associated with DNA-repair deficiency and commonly mutated in prostate cancer were targeted for mutations and deletions (Supplementary table 1). Accompanying sequencing of germline DNA revealed high-impact variants in 8.92% (*ATM, BRCA1, BRCA2* and *CHEK2*), similar to recent reports [56–58]. However, only 2/213 (excluding four germline DNA samples that failed processing) carried pathogenic *BRCA2* mutations, significantly less than Pritchard et al [57] and Mandleker et al [58] (Fisher's exact test: p = 0.00328, p = 0.000735, respectively). Both reported multiple occurrences of Ashkenazi Jewish founder mutations such as the *BRCA2* p.Ser1982Argfs*22, not observed in this report, suggesting a difference in the underlying population demographics (Supplementary table 5). Excluding MSI positive cases and focusing on somatic clonal high impact variants, homozygous deletions or germline variants in DNA-repair genes, 18 (8.29%) individuals had biallelic inactivation whereas 39 (18.0%) had one detectable alteration (Figure 5, Supplementary table 4). Note, however that the intronic regions were not targeted in the current version of these capture designs rendering structural variation undetectable, except close to exons or baits designed for CNA purposes.

**Figure 5.**
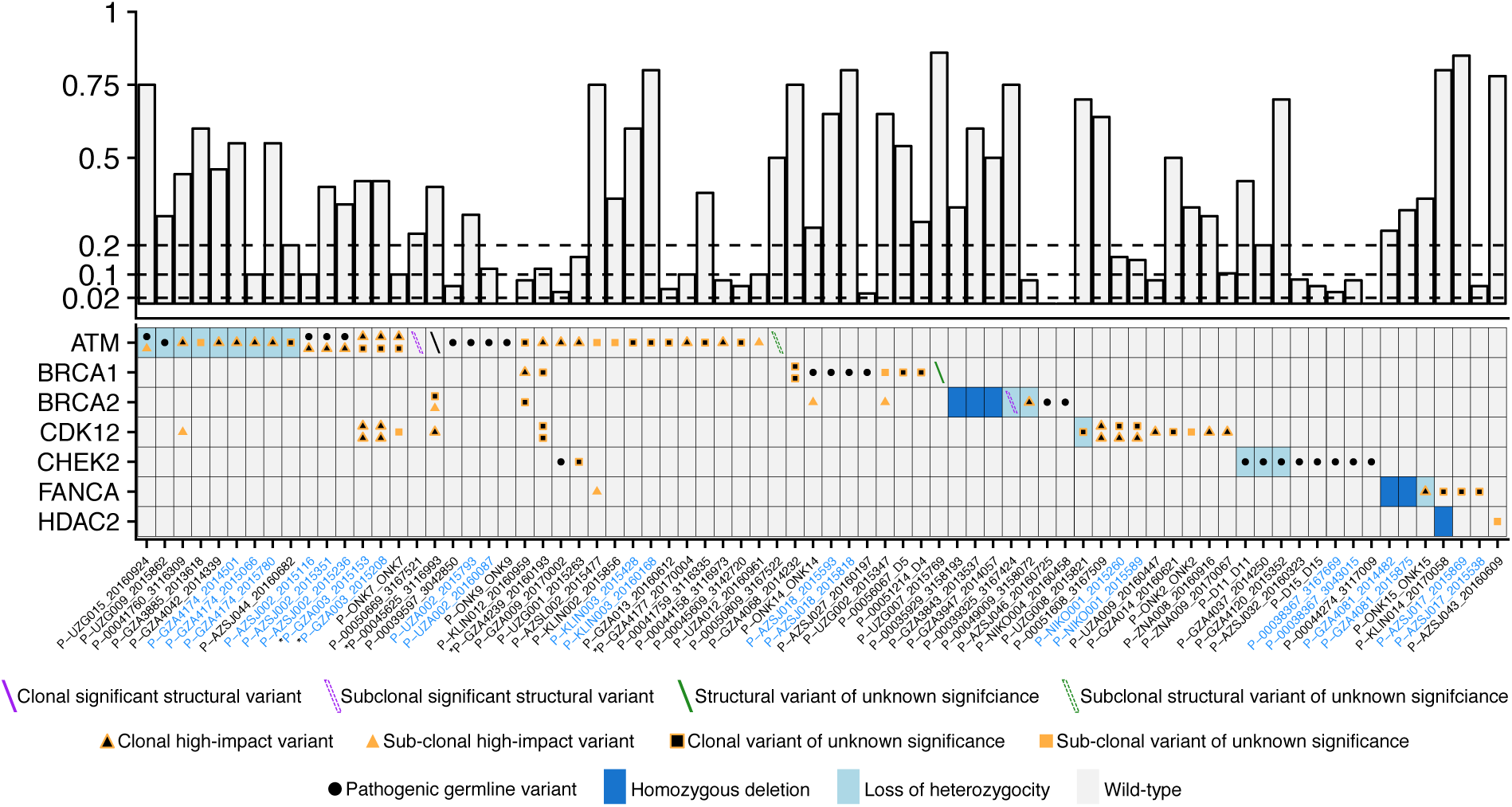
Alterations in genes associated with DNA repair deficiency. The upper panel displays the circulating tumor DNA fraction. The dashed lines at 0.02, 0.10 and 0.20 denotes the cutoff to detect point mutations, loss of heterozygosity and homozygous deletions, respectively. Bottom panel, heatmap of the mutational landscape detected from circulating tumor DNA profiling. Type of alteration is coded according to the bottom legend. For visualization purposes, maximally two mutations or structural variants are displayed per patient. Triangles and boxes represents single nucleotide variants and indels. Subclonal mutations are defined as having an allele frequency < 1/4 of the ctDNA fraction. The same definition was applied to structural variants after median allele allele frequency adjustment versus the mutations (Methods). The *BRCA2* structural variant of patient P–00039325, sample 3167424 was classified as borderline subclonal although relevant in the progressing clone after chemohormonal treatment (Supplementary figure 6C). Synonymous point mutations are not displayed here. Variants of unknown significance are non-synonymous single nucleotide variants outside hotspots, not annotated as pathogenic from variant databases. Structural variants of unknown significance are e.g. confined to a single intron, without affecting neighbouring exons. X-axis: Cell-free DNA samples sorted according to the number of alterations detected in each gene in alphabetical order. Patients with multiple samples are colored in blue. * samples with microsatellite instability.

### Clonal hematopoiesis causes false positive findings

Aberrant blood cell populations [37–40] have the potential to confound ctDNA mutational profiles when performed without matched blood DNA as a control. To assess the potential impact and prevalence of genetically aberrant blood cell expansions in our cohort, we investigated copy number and mutational data for indications of aberrations present in both cfDNA and WBC DNA. We observed, in separate patients, four cases of large arm-level copy number alteration (chr 11,13 and 20) in white blood cells with coverage ratio and SNP allele ratio suggesting a cellularity between 40–65%, and a focal CCND1 amplification with coverage ratio 1.57, similarly detected in the cfDNA. Putative somatic point mutations were interrogated in white blood cell DNA using pooled healthy donor DNA as control and excluding variants exceeding 25% allele ratio, outside hotspots, as likely germline. Thirty seven protein altering variants were observed in another 29 patients and could be validated in patient-matched cfDNA, including hotspot mutations in *AKT1, BRAF, CTNNB1, DNMT3A, NRAS, SF3B1* and *TP53* (Figure 6). In summary, 40 false positive variants in 31 patients (14.6%) would have been included in ctDNA mutational profiles if matched white blood cells had not also been sequenced.

**Figure 6.**
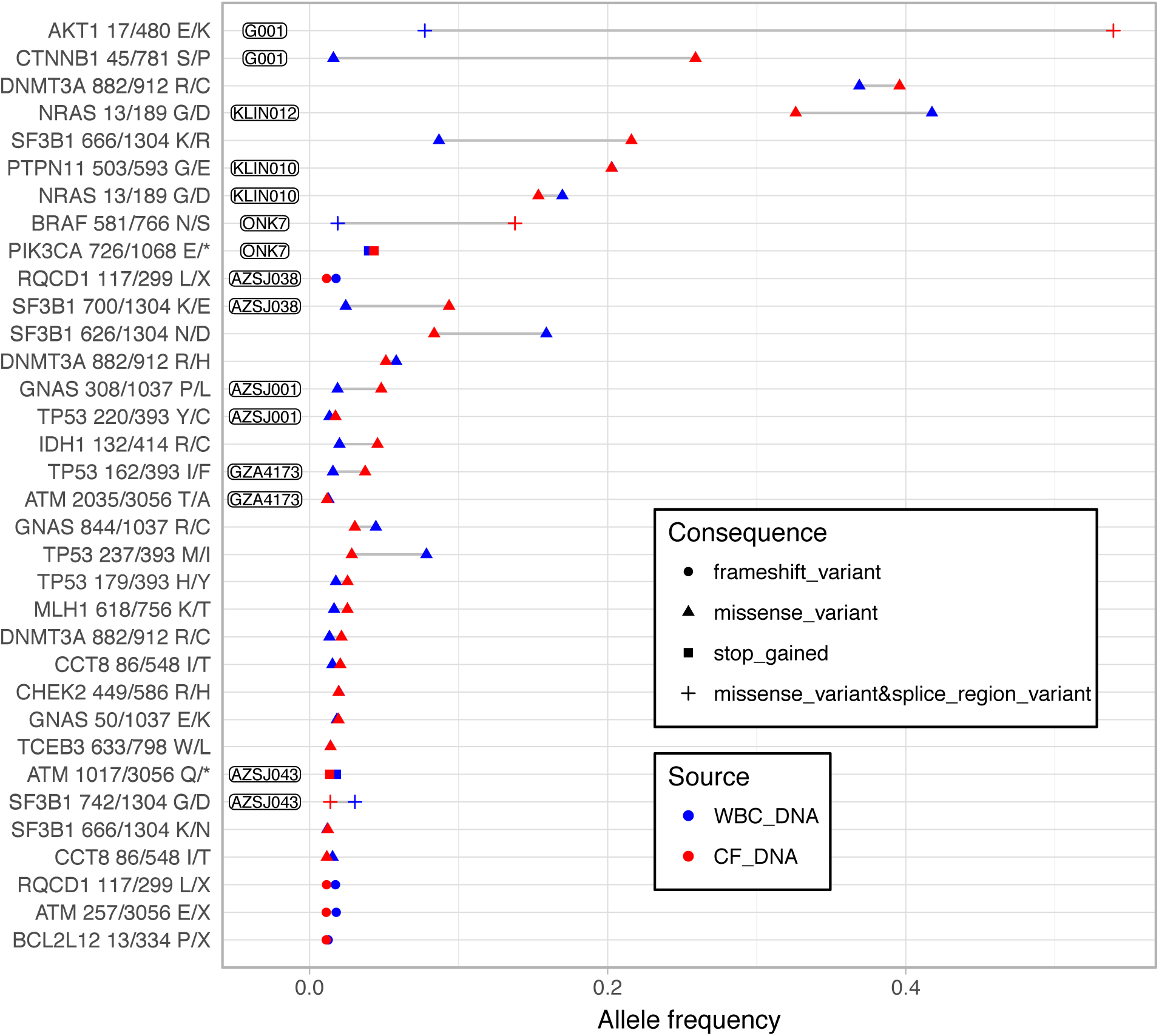
Clonal hematopoiesis. Somatic point mutations detected in germline DNA extracted from white blood cells and validated in cell-free DNA. For each point mutation, the amino acid position and total number of amino acids are given. Patients with multiple mutations are labeled with patient ID. X-axis; Variant allele frequency. Y-axis; individual point mutations sorted according to allele frequency in white blood cells and individual. The inset legend explains the type and the source of each variant.

## Discussion

Genomics-guided therapy selection is arguably the most promising avenue to remedy trial-and-error treatment decisions and the accelerating costs of drugs [21]. However, the utility of tumor profiling is currently limited in mPC due to the lack of validated predictive biomarkers. Liquid biopsies have the potential to act as a tissue substitute and cost-efficiently accelerate trials designed to identify predictive biomarkers. Therefore, we set out to comprehensively profile cfDNA samples in mPC, encompassing mHNPC to mCRPC, to gain knowledge relevant for applying ctDNA in a clinical trial context. Our key findings are: 1) ctDNA fractions increased gradually from first- to fourth line of therapy. Baseline samples had higher ctDNA fraction compared to follow-up samples, but the difference became non-significant after the second line of mCRPC therapy; 2) biallelic inactivation of key tumor suppressors always occurred in high tumor burden samples, with only one exception, providing a rationale for low ctDNA fraction samples with poor sensitivity to detect the second hit; 3) clonal high-impact structural variation is twice as common as point mutations, which challenges the traditional focus on coding regions; 4) the three potentially predictive biomarkers in mPC, microsatellite instability, mutations in genes associated with DNA-repair deficiency and *AR* aberrations were detected at expected rates; 5) clonal hematopoiesis occurs frequently, demanding synchronous WBC profiling to avoid false positive genotyping.

Due to the genomic diversity of metastatic cancer, resistance will always arise to single agent therapies where the duration of response is correlated to the number of cancer cells in a patient [59]. Towards end stage disease, progression will occur more rapidly, regardless of therapy, with the exception of extreme responders to immunomodulators [14]. Molecular biomarker-driven clinical trials are commonly targeting patients where no approved treatment options remain, although primary outcomes may be hard to achieve if disease burden is too high [60]. Paradoxically, we find that liquid biopsies carry more information towards end stage disease, and are currently of limited value in a significant fraction of patients starting first- and second line mCRPC therapy due to low tumor burden (Figure 1). A potential solution may be a complementary approach using CTCs to gain insight into ploidy and CNA and ctDNA for mutations and structural rearrangements. However, there are some limitations: we show that CTC counts correlate with ctDNA fraction (Supplementary figure 1), and patients with low ctDNA fraction starting first- and second line mCRPC therapy, with few exceptions, have low CTC counts (Supplementary figure 9); previous work demonstrate poor success rate (~10 %) in obtaining high-quality CTC sequencing data from isolated cells [61,62] necessitating multiple 10 ml blood tubes for CTC analysis in first- and second line patients. However, recent improvements in harvesting metastatic tissue [29] may provide a fall-back if ctDNA profiling fail to identify any relevant biomarkers. As the success-rate of harvesting high-quality metastatic tissue is also correlated to tumor burden [26,27], prospective validation is needed to establish the most feasible approach.

The inherent challenges to complement ctDNA profiling inspired us to investigate the necessity of a second hit to infer tumor suppressor deficiency. By deep sequencing of all non-repetitive intronic and exonic regions in *TP53, PTEN* and *RB1* in high ctDNA fraction samples, we investigated if the detection of one clonal high-impact variant is adequate to infer biallelic inactivation. Out of 71 samples in 59 men with ≥0.2 ctDNA fraction, 47.5%, 20.3% and 44.1% harbored biallelic inactivation of *PTEN, RB1* and *TP53*, respectively (Figure 3). Only one patient carried a clonal high impact variant, a deletion in *TP53*, without a detectable event on the other allele. These data are encouraging as a large fraction of *TP53* was not possible to sequence due to repetitive DNA (Supplementary figure 7). The observation is consistent with exome sequencing of 150 mCRPC tissues which revealed that biallelic inactivation always occured if one high-impact event was observed in a key tumor suppressor such as *PTEN* or *RB1* [7]. Interestingly, residual break-points remained in 5/17 samples with a homozygous deletion in *PTEN*, which is detectable, even when tumor burden is low.

Two commercial alternatives for ctDNA-based profiling were recently compared with surprisingly low concordance [63]. The lack of accompanying white blood cell germline profiling makes it hard to separate germline variation from somatic [64] and impossible to distinguish clonal hematopoiesis [37–40] from ctDNA. In our study, 14.6% of patients harbored clonal expansions in the WBC compartment. However, the majority of men with mCRPC probably suffer from clonal hematopoiesis as the targeted sequencing applied here only covered 60 out of 327 driver mutations associated with clonal expansions in the blood [38]. Taken together with our findings, we strongly discourage the use of commercial assays that only analyse cfDNA from plasma.

We demonstrate that ctDNA can be applied to comprehensively profile *AR* and identify alterations in genes associated with DNA repair deficiency. Although these two biomarkers are considered promising for clinical decision making, recent data question the association between DNA-repair germline variants and response to PARP inhibitors [65] and the independent association between *AR* amplifications and response to abiraterone and enzalutamide treatment in mCRPC [55]. In the light of the controversy over AR-V7 [24], the only possible conclusion is the urgent need for efficient biomarker-driven clinical trials to identify in detail which patients may benefit from each therapy option.

We present two major novel findings: firstly, we show that the MSI phenotype may be detected directly from cell-free DNA. We believe that this novel findings will be impactful for the prostate cancer community in the light of the recent approval of pembrolizumab for MSI-positive solid tumors [11]. Secondly, we demonstrate that high-impact structural variation is approximately twice as common as high-impact point mutations in *PTEN* and *RB1*, which questions which challenges the traditional focus on coding regions in the context of clinical trials and clinical decision making.

## Conclusions

This study strengthen the accumulating evidence that ctDNA profiling mirrors the somatic alteration landscape from metastatic tissue by demonstrating, for the first time, that the MSI phenotype may be detected directly from cell-free DNA. However, to enable acceleration of clinical trials through ctDNA analysis, intronic sequencing of tumor suppressors in combination with synchronous profiling of white blood cells must be applied to prevent inaccurate somatic alteration assignment, which in turn, may reduce the power to identify predictive biomarkers.

## Authors’ contributions

BDL, PR, JL and HG conceived the study and designed the methodologies and experiments. BDL, SO, JL and GVdE performed experiments and acquired data. BDL, MM, TW, JL and MR analysed and interpreted the data. PVO, CG, JA, PO, WD, LH, DS, BB, WL, EE, DDM, MS, KF, AR, AU, JY, HG, TN, MA, PR and LD recruited patients and acquired blood samples for analysis. BDL, JL, MM and HG prepared the manuscript. BDL, SO, MM, LE, TW, PJvD, PVO, CG, JA, PO, WD, LH, DS, BB, WL, EE, DDM, MS, KF, AB, DG, LH, GVdE, AR, MR, PR, AU, JY, HG, SVL, JL and LD performed critical revision of the manuscript for important intellectual content.

## Acknowledgements

The Belgian Foundation Against Cancer (grant number C/2014/227); Kom op tegen Kanker (Stand up to Cancer), the Flemish cancer society (grant number 00000000116000000206); The Cancer Research Funds of Radiumhemmet, through the PCM program at KI (grant number 163012); The Erling-Persson family foundation (grant number 4-2689-2016; the Swedish Research Council (grant number K2010-70X-20430-04-3) and the Swedish Cancer Foundation (grant number 09-0677). This funding bodies had no role in the design, execution or interpretation of the data in this study and did not influence the decision to submit results for publication. The authors thank all patients for their willingness to participate in this study. We also thank Luc De Laere, Thijs Develter, Sophie Vantieghem, Sofie Herman, Gwen Colfs, Veerle Lamotte, Anita Boumans, Abdelbari Baitar, Roos Haeck, Goele Wallays, Hanna Emanuelsson, Per Gustafsson, Jenny Wängberg, Maria Bergh and Claudia Maes for their assistance with patient inclusion, sampling management and data collection.

